# A leaf phenomics approach to estimating below-ground traits in North American Licorice

**DOI:** 10.1101/2025.04.07.647639

**Authors:** Zachary N. Harris, Vivian Tran, Emelyn Piotter, Meredith T. Hanlon, Matthew J. Rubin, Allison J. Miller

## Abstract

**Premise of the study:** Thousands of years of selective breeding has prioritized above-ground yield, with little regard for changes happening below-ground. Despite their central role in plant success and resilience, our knowledge of roots lags behind above-ground structures. Accurately phenotyping root traits is often labor-intensive, expensive, and destructive. In order to advance understanding of the fundamental biology underlying root systems, and to integrate hard-to-measure root traits into breeding programs, high-throughput non-destructive methods are required.

**Methods:** This study uses American licorice (*Glycyrrhiza lepidota* Pursh.), a perennial legume with a rich ethnobotanical history, as a model to investigate root system phenotypes. We assess root traits across multiple populations, analyze relationships between above- and below-ground phenotypes, and test the use of multidimensional leaf traits, including spectral reflectance, in predicting root traits.

**Key results:** American licorice displays significant variation in root traits across source populations and strong correlations between above- and below-ground traits. Leaf spectral reflectance and elemental composition show promise in modeling below-ground traits, though the isometric relationship between plant size and root traits complicates model accuracy.

**Conclusions:** These findings demonstrate the use of high-dimensional leaf traits as a proxy for root traits, with potential applications for understanding foundational questions in plant biology and in breeding programs targeting the below-ground structures of perennial herbaceous species. Further optimization and larger studies are needed to improve predictive models.

## INTRODUCTION

Thousands of years of crop domestication have altered root system phenotypes, primarily as a result of selective breeding focused on above-ground yield (Martín-Robles et al., 2019). While some crops have been selectively bred for below-ground traits, such as edible roots or tubers, even these systems have experienced trade-offs between edible and non-edible tissue investment (Denham et al., 2020). In maize, one of the most extensively studied crops, advances in both breeding and management for increased yield have also altered root traits. Modern maize, which has been bred to grow at high planting density, has decreased root system length and biomass (Rinehart et al., 2024) and increased plasticity for improved nutrient acquisition (York et al., 2015; Sciarresi et al., 2025). Similar trends across annual crops illustrate how prioritizing above-ground traits often leads to altered root systems, some of which have reduced the plant’s ability to maintain soil health, mitigate erosion, and support nutrient cycling (Zhang et al., 2011; Smaje, 2015; Crews et al., 2018). Modern agricultural practices, such as generous use of fertilizers and soil manipulation, further exacerbate these issues. Future agricultural challenges will require that root systems are more efficient at nutrient and water capture while also slowing, or even reversing, land degradation (Lynch, 2022; Lynch et al., 2022; Mehra et al., 2025). Addressing these needs requires a better understanding of root systems and their contributions to both plant productivity and ecosystem services (Griffiths et al., 2022).

Accurate and high-throughput phenotyping of root traits remains a key limitation in plant biology, ecology, and breeding research programs. Root systems show variation in morphology, network topology, distribution of roots and root mass in space, and whole system architecture (Lynch, 1995; Seethepalli et al., 2021), but even simple phenotypes like rooting depth are exceptionally difficult to measure in field conditions at scale (Hanlon et al., 2024). Under controlled conditions, phenotypes like root depth can be measured but often under experimental conditions that do not capture those experienced by plants in nature, like vertical agar plates (Ogura et al., 2019) or hydroponic systems (Mathieu et al., 2015). In the field, labor intensive techniques for studying roots include destructive sampling (excavation), soil coring, minirhizotrons, imaging with large instrumentation, and the use of tracers (Cabal et al., 2021; Hanlon et al., 2024). Integrating root system phenotyping into large-scale experiments or high-throughput breeding programs remains a challenge because phenotyping belowground organs is usually destructive (Kuijken et al., 2015), which is especially problematic for species that persist for multiple years. If methods could be developed for rapid, non-destructive characterization of root traits, belowground plant structures could be more easily integrated into basic plant biology research and could become more reasonable targets for breeding (Maqbool et al., 2022).

If root systems are to be more intensively studied or targeted in breeding programs, improved methods for phenotyping below-ground traits are required. Multidimensional phenotyping methods have shown promise for modeling shoot system traits that are otherwise difficult and/or expensive to measure. One of the most active areas of research uses above-ground (usually leaf or canopy) spectral reflectance to predict other plant traits (Moreira et al., 2020; Kothari and Schweiger, 2022). Models based on hyperspectral data have been used to predict disease presence (Yu et al., 2018), metabolic composition (Vergara-Diaz et al., 2020), and plant developmental stage (Gutiérrez et al., 2019; Kaur et al., 2024), among others. Hyperspectral models have also been developed for root traits using hyperspectral reflectance from the roots themselves (Bodner et al., 2018; Chang et al., 2024; Faehn et al., 2024). Further, work has suggested that above-ground phenotyping modalities like leaf elemental composition can predict root length (Hanlon et al., 2024). The goal of this study is to test the utility of above-ground, multidimensional data (spectral reflectance and elemental composition) measured in shoots as a proxy for root phenotypes in a perennial herbaceous system.

Although perennial, herbaceous crops were not developed by early farmers (Van Tassel et al., 2010), they are garnering increasing attention as promising emerging crops that produce economically viable products and offer important ecosystem services (e.g., (Zhang et al., 2022)). The roots of many perennial, herbaceous species have been used historically for both food and medicine. These species exhibit extensive, quantifiable variation in root traits and offer a valuable system for understanding how multidimensional above-ground phenotypes relate to below-ground phenotypic variation. Among these, species within the licorice genus (*Glycyrrhiza* L.) have been grown by peoples in different parts of the world for centuries (Kindscher, 1992; Tang and Eisenbrand, 1992; Richardson, 2002). One of the most commonly cultivated licorice species, *Glycyrrhiza glabra* L., is native to southern and eastern Europe, middle and western Asia, and northern Africa (GRIN-global, 2025). *Glycyrrhiza glabra* is cultivated for food additives (mainly as a sweetener (Richardson, 2002)), dermatology and cosmetics (Burlando et al., 2010), and pharmacology (Tang and Eisenbrand, 1992; GRIN-global, 2025). A related species, American licorice (*Glycyrriza lepidota* Pursh.), native to the North American Great Plains, has a similarly rich ethnobotanical history (Kindscher, 1992; Moerman, 1998, 2009; Paul J. Johnson, Abigail P. Martens, Arvid Boe, 2023). Both *G. glabra* and *G. lepidota* are cultivated for compounds found in their root systems, some of which are directly correlated with root phenotypes like biomass and diameter (Hou et al., 2018). Rapid, non-destructive root phenotyping platforms could be immensely valuable for the future study and optimization of systems like licorice, which persist for multiple years and are valued for below-ground structures.

This study aims to address critical gaps in phenotyping root systems using American licorice as a model. Specifically, we investigate: 1) differences in phenotypes (above- and below-ground) as a function of plant size and source populations, 2) correlations within and among above-ground and below-ground phenotypes, and 3) the potential of high-dimensional leaf traits to model below-ground traits. In this study, we focus on below-ground traits that are common in all plant species (e.g., root biomass, length, diameter, and surface area). We find that root traits show variation based on source population and exhibit strong relationships with above-ground biomass. We formalize these relationships using allometric models for each measured trait. We then show the capacity of high-dimensional above-ground phenotypes to predict below-ground root traits with and without accounting for above-ground biomass. This work suggests both spectral and elemental models derived from leaves are promising avenues of future research for foundational plant biology and for advancing below-ground targets of selection in breeding programs.

## MATERIALS AND METHODS

Seeds from 19 *G. lepidota* populations representing seven states were obtained from the Chicago Botanic Garden (3 populations), United States Department of Agriculture Germplasm Resources Information Network (USDA GRIN; 9 populations), Dr. Arvid Boe (4 populations), and commercial sources: Everwilde Farms, Prairie Moon and Sheffield’s Seeds (Appendix S1). On May 5, 2021, 12 seeds from each population were planted into a 96-cell propagation flat with Berger Bm7 35% Bark HP soil (Saint-Modeste, Quebec, Canada) and placed into a growth chamber at the Danforth Plant Science Center (St. Louis, MO, USA) with the following conditions: 16h day length (200 μmol•s^-1^•m^-2^), 22°C, and 50% humidity. Germination was recorded daily. At three weeks post-germination, a total of 78 germinants representing 18 populations were transplanted into 3.7 L pots with Berger Bm7 35% Bark HP + 14-14-14 Osmocote (14 g/pot) media and moved to an outdoor nursery at the Danforth Center. While in the nursery, plants were watered as needed.

### Shoot system phenotyping

#### Leaf spectral data

After 5 months of growth (October 2021; mean germination time ∼4 days), we collected three fully-expanded leaves from each plant for multidimensional shoot system phenotyping (n=72 plants, 3 leaves per plant; Figure 1A). The sampled leaves were imaged using the CropReporter (PhenoVation, Wageningen, NL) which measured broadband reflectance in the blue (∼470 nm), green (∼530 nm), SpcGreen (∼550 nm), red (∼660 nm) and Near-Infrared (NIR; ∼790 nm) portions of the spectrum. In addition, the CropReporter reports average hue, saturation, value, F0, Fm, dChl, and dFv/Fm. From these images, we extracted maximum quantum yield of photosystem II (Fv/Fm), Normalized Difference Vegetation Index (NDVI), chlorophyll index, and anthocyanin index. For simplicity, all crop reporter traits were reduced via principal component analysis (PCA) to retain ∼75% of the total CropReporter variation (cPCs 1-3; Appendix S2).

**Figure 1:**
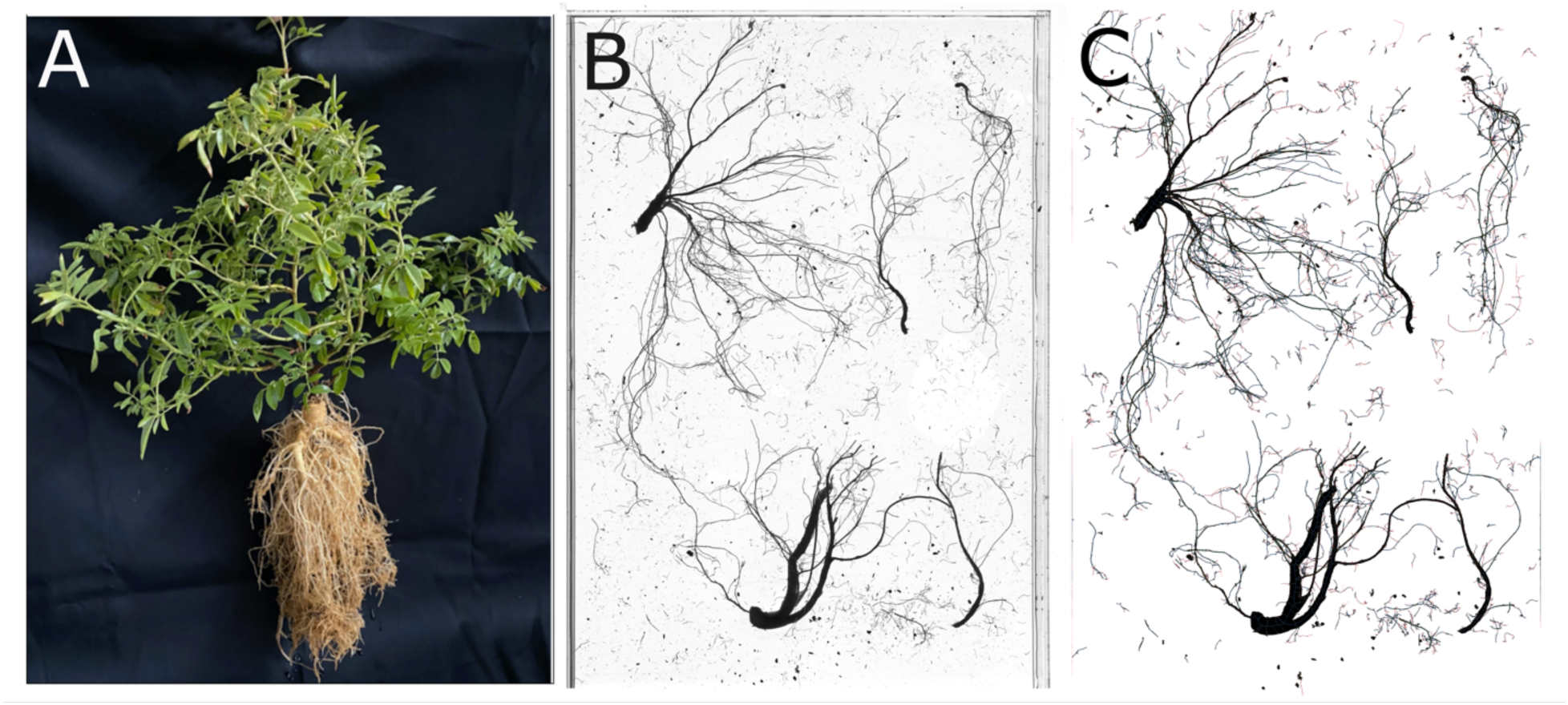
Study organism and visualization of root phenotyping. (A) Image of a *Glycchyriza lepidota* sample after harvest for phenotyping and potting media had been removed from the root system. (B) A single example root system scan prior to analysis. C) Cleaned and segmented root scan following processing with RhizoVision Explorer.

Spectral reflectance of leaves was captured using a LinkSquare 1 (LinkSquare, San Jose, CA, USA), which measures reflectance between 400 and 1050 nm. The Linksquare 1 contains two light sources: LS0, optimized for measurement in the visible range to red-edge, and LS1, optimized for measurement in the red-edge to NIR range. The LinkSquare was placed on four locations on the adaxial leaf surface of four fully expanded leaflets pressed to a constant black fabric wrapped piece of cardboard as a backing. The LinkSquare was paired via Bluetooth to an Android tablet running the app Prospector (Rife et al., 2021). Outlier scans were detected and removed using the R package ‘waves’ (Hershberger et al., 2021) based on the distribution of Mahalanobis distances calculated between each sample and the total distribution of spectra, filtering at a 95% cutoff based on a chi-square distribution parameterized with degrees of freedom equal to the number of wavelengths. We then tested nine spectral pretreatments using ‘waves’ across all root phenotypes (described below) to understand which pretreatment would likely produce the best models for root phenotypes. The 9 pretreatments were: raw reflectance, reflectance through a Savitsky-Golay filter (SG, w=11), a standard normal variate filter (SNV), the first derivative of the first three treatments, and the second derivative of each of the first three treatments (Appendix S3). On average, it took 31 PCs to explain ∼75% of the spectral variation across each pretreatment. Multiple regression models were computed for each pretreatment for each phenotype and the pretreatment that produced the highest average R-squared values (SNV) was selected. We opted to keep all 31 SNV-pretreated spectral PCs despite the top 31 PCs explaining ∼81% of total variance as opposed to 75% for the other modalities, as reducing the number selected yielded a different ‘best’ pretreatment (Appendix S4).

#### Leaf elemental concentrations

We sampled 204 total leaves (68 plants x three leaves per plant = 204) for analysis of elemental composition. Each leaf was dried in an oven at 60°C, ground, and subsampled to a weight between 75 and 100 mg, where possible. Samples were analyzed using an established pipeline at the Donald Danforth Plant Science Center (Ziegler et al., 2013) using ICP-MS which simultaneously quantifies the concentration of 20 ions: calcium (Ca), cobalt (Co), copper (Cu), iron (Fe), magnesium (Mg), manganese (Mn), molybdenum (Mo), phosphorus (P), potassium (K), selenium (Se), sodium (Na), sulfur (S), zinc (Zn), aluminum (Al), arsenic (As), boron (B), cadmium (Cd), nickel (Ni), rubidium (Rb), and strontium (Sr). Samples with <20mg of tissue were excluded from further analysis and each element was averaged across the three replicates taken from each plant. As above, elemental PCs were selected such that ∼75% of elemental variation was captured (ePCs 1-5; Appendix S5).

#### Shoot biomass

We harvested above-ground biomass of the plant. Samples were dried at 65C for three days and weighed.

### Root system phenotyping

While human use of licorice roots targets chemical properties, here we focus more broadly on root traits that are relevant for, and can be quantified in, a broad diversity of species. We selected these root traits as a first-pass, proof of concept effort to test whether above-ground features can predict below-ground structures.

Below-ground biomass was harvested at the same time as above-ground biomass. Planting media was removed from roots through soaking and spraying. Cleaned root samples were stored in plastic bags at -20°C prior to scanning. After thawing, remaining planting media was washed from samples and root systems were placed in a clear tray with water where roots were separated from one another. The clear tray was placed on a flatbed scanner (Epson 12000XL), and top-down scans were obtained using the transparency unit at 1200 dpi and saved as high-quality JPEG files using the Epson Scan 2 software. Following scanning, roots were dried at 65°C for three days and weighed to the nearest 0.01 gram (below-ground biomass). The ratio of above- and below-ground biomass was computed as an additional phenotype.

Root systems scans were segmented and features were extracted using Rhizovision Explorer with the following deviations from default parameters: root type = Broken roots, maximum background noisy component size = 2, enable edge smoothing = true, enable root pruning = true, root pruning threshold = 20 (Figure 1B-C) (Seethepalli et al., 2021). From each scan, the image resolution was used to convert from pixels to mm. Diameter range classes were set to 0.25 mm increments until diameter range class 11 (the Rhizovision maximum), which was composed of all diameters > 2.5 mm. Because root system scans were taken across several images, root traits were combined using a RhizoVision Explorer companion script (Seethepalli et al., 2024). Trait distributions suggested that 11 diameter range classes were unnecessary, so all classes beyond diameter class 6 were concatenated into DR6+ (>1.25 mm). The total list of root traits was reduced to remove traits that were inferred in 3D (volume and volume in all diameter classes), traits that were deemed inaccurate due to how the root system was broken up on the scanner (perimeter, number of root tips, branching points, and branching frequency), and traits that were known to be nearly exactly correlated to other traits (projected area and network area, which are nearly perfectly correlated with surface area). After this reduction, the following root system traits were analyzed: total root length, average root diameter, maximum root diameter, root system surface area, root length contained in each of the six diameter classes, and surface area contained in each of the six diameter classes. In addition, we computed specific root length (SRL, the ratio of total root length and below-ground biomass), the proportion of root length contained in each of the six diameter classes, and the proportion of surface area contained in each of the diameter classes.

### Tests for plant size allometry

One of the key challenges in predicting below-ground traits using above-ground phenotypes is untangling variation that is unique to root traits and is not merely a reflection of plant size. We tested for root trait isometry against four different measures of plant size. First, we regressed log transformed root traits against log transformed above-ground biomass. Second, we regressed log transformed root traits against log transformed below-ground biomass. Third, we fit a linear model regressing above-ground biomass against below-ground biomass and used the log-transformed residuals from this model to represent variation in above-ground biomass that was unique to above-ground biomass, in other words, that excluded variation in below-ground biomass. Finally, we did the same thing for below ground variation that excluded variation in above-ground biomass. In all cases, the slope of the line was evaluated for a value near one which would indicate isometry. Values near zero were taken to represent no relationship. Intermediate values represented allometric relationships.

### Population effects, trait correlations, and bivariate spectral models

Fixed effects linear models were used to understand the influence of population on above- and below-ground traits, adjusting for above-ground biomass (included as a non-interacting, fixed effect). Bivariate correlations were calculated using the Hmisc (Harrell, 2022) and corrplot (Wei and Simko, 2021) packages. P-value correction for multiple bivariate trait correlations was not applied because the traits were expected to exhibit a high degree of correlation (for example, root traits within and across size classes) making multiple test correction overly conservative and likely obscuring biologically meaningful relationships. Linear models regressing traits against wavelength-by-wavelength spectral reflectance were parameterized with scaled root traits being modeled as a function of scaled reflectance (following the SNV spectral pretreatment) at a particular wavelength.

### Multivariate models

Multivariate principal component regressions were parameterized with scaled root traits being modeled as a function of the top ∼75% of variation captured by CropReporter (3), spectral (31), or elemental (5) PCs, as fixed, additive, non-interacting effects. Models were fit using a leave-one-out cross validation paradigm where (N-1) models were fit and then used to predict the sample left out for each fold. Goodness of fit for these models was assessed by regressing the true value for each trait against the predicted value from the CV fold it was withheld from and computing the R^2^ from that regression. Models were fit by regressing each trait as a scaled value and after adjustment for above-ground biomass. All analyses presented in this manuscript were completed in R v4.2.1 (R Core Team, 2022).

## RESULTS

### Variation in below-ground traits across source populations

In this study, we quantified variation in above-ground biomass and 31 root-associated traits, including below-ground biomass, the ratio of above- to below-ground biomass, specific root length, total root length, average root diameter, maximum diameter, and root system surface area. Two traits (total root length and root system surface area) were analyzed in six diameter classes from 0 to 1.50+ mm in 0.25 increments. The total range of variation for each trait, including the minimum value, 25% quartile, median value, mean value, 75% quartile, and maximum value are reported in Appendix S6.

To account for potential differences in overall plant size, we modeled each below-ground phenotype while controlling for above-ground biomass. As expected, most below-ground traits measured in this study were significantly associated with above-ground biomass (Figure 2A). Nearly all root traits, including total root length and surface area, were associated with larger above-ground biomass, with above-ground biomass explaining between 7.2 and 32.7% of total root trait variation. However, certain traits, such as the ratio of above- to below-ground biomass, specific root length, and the proportion of root length and surface area in diameter range classes 2 and 5, did not show this association, suggesting that plant size alone does not explain variation in these traits.

**Figure 2:**
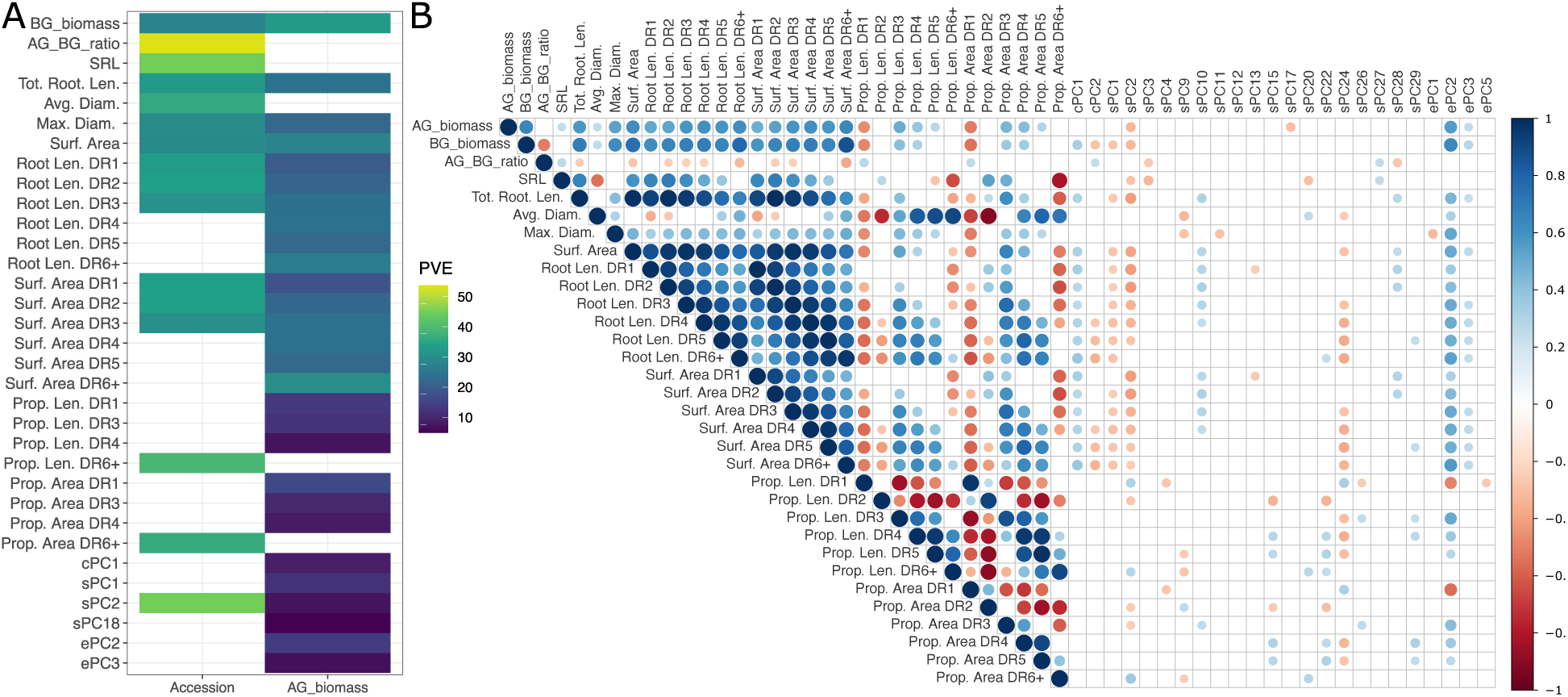
Source population influence and bivariate trait correlations. (A) Heat map showing the variance explained in each trait by the source population and above-ground biomass. Traits excluded from this figure were not significant for source population or above-ground biomass. (B): Correlation matrix of all traits in this study. Pearson correlation coefficients are reflected in both the size and hue of the circle for each pair or traits. Above-ground PC-based traits missing from this figure showed no significant correlations with any other trait. The rest of the correlation matrix, representing only the correlations within above-ground traits, can be found in Appendix S8.

In addition to plant size, we observed significant variation in several root traits across source populations (Figure 2A). The strongest effect from source population was observed in the ratio of above- to below-ground biomass. Other traits influenced by source population were below-ground biomass, specific root length, total root length, and surface area. Root length and surface area within diameter range classes 1-3 were variable by population, and the proportion of root length and root area in diameter class 6+ was different across source populations. When significant, source population explained between 28.0 and 53.6% of total root trait variation.

### Generation and analysis of multidimensional above-ground phenotypic data

To investigate capacity of above-ground structures to predict below-ground traits, we reduced the dimensionality in each of the three modalities of above-ground predictors. In doing so, we generated three sets of PCA-derived predictor variables: cPCs (derived from data generated by the CropReporter), sPCs (spectral data derived from the LinkSquare), and ePCs (elemental composition data). To explain the composition of these PCs, we looked at the variation explained by and the features loading most strongly onto each PC. The first 3 cPCs explained ∼75% of the CropReporter data. cPC1 (44.4%; Appendix S2) was most heavily influenced by green reflectance (positive) and chlorophyll and anthocyanin indices (negative). cPC2 (24.3%) was most heavily influenced by broadband NIR reflectance (positive) and saturation (negative). cPC3 (8.8%) was most heavily influenced by far red reflectance (positive) and dCHL (negative). For the spectral data, the top three sPCs explained ∼51.8% of the observed variation (37.9%, 8.8% and 5.1% explained by PC1, PC2, and PC3, respectively). Wavelength-by-wavelength loadings across both sensors in the LinkSquare are visualized in Appendix S7. For the elemental composition data, the top three ePCs explained 61.7% of the variation (Appendix S5). All elements loaded negatively onto ePC1 (44.0%), with the heaviest loaders being sulfur and boron. Potassium and phosphorus dominated the negative loadings on ePC2 (10.2%) with nickel and strontium loading most positively. ePC3 (7.5%) was most influenced by molybdenum and nickel (positive) and arsenic (negative). ePC4 (6.1%) had cobalt as the strongest positive loading element and sodium and the strongest weakest loading element. Finally, ePC5 (5.2%) was most influenced by copper (positive) and zinc (negative).

Each of the principal component predictors was also assessed for variation due to above-ground biomass and source population. Interestingly, above-ground predictor traits were generally less influenced by either above-ground biomass or source population than root traits. Notable exceptions include cPC1, sPC1-2, sPC18, and ePCs 2-3, which were associated with above-ground biomass (4.9 - 13.7% variation explained). Among these, only sPC2 showed significant variation by population (44.9% variation explained) (Figure 2A).

### Trait correlations

We next tested the correlation structure between all possible pairs of traits (Figure 2B). Out of 1431 possible correlations, we found that 570 (39.8%) were significant (P < 0.05). Of these, 378 (66.3%) were positively correlated and 192 (33.7%) were negatively correlated. The vast majority of correlations identified in the study are either within below-ground traits or across below- and above-ground traits. Very few correlations within above-ground traits were identified in this study (Appendix S8).

Generally, we found a large suite of root traits correlated positively amongst themselves. This suite included below-ground biomass, specific root length, total root length, maximum diameter, surface area, root length in all six diameter range classes, and surface area in all diameter range classes. For example, total root length was positively correlated with the total root surface area (r = 0.98, P < 0.01) and maximum diameter was correlated with surface area (r = 0.46, P < 0.01). Notably missing from this cohort was the ratio of above- and below-ground biomass and average root diameter which were both negatively associated with different phenotypes in the large root suite. Within root traits, other negative correlations were primarily identified in traits representing proportions. For example, the proportion of root length in diameter class 2 (0.25 - 0.50 mm) was negatively associated with the root length contained in higher diameter range classes. Both the proportion of root length and surface area in diameter range class 6+ (> 1.50 mm) was negatively associated with root length and surface area in most diameter classes. This suggested that there was a trade off in the root system resource allocation - more allocation to tap roots was associated with less allocation in smaller diameter classes.

Every root phenotype measured in this study was associated with at least one above-ground predictor trait (PCs from CropReporter, spectral, and/or elemental data). For example, sPC1 and sPC2 were negatively associated with the large suite of correlated root traits while sPC10 and ePC2 were positively associated with that suite of traits. Subtle but interesting relationships include SRL, which was correlated with sPC2, 3, and 20 and average root diameter which was correlated with sPC9, 20, and 24.

### Trait allometry

To determine whether variation in root traits was distinct from overall plant size we regressed log transformed root traits against four log-transformed components of plant size (see Methods). We found that total root length, surface area, root length in five of the six diameter range classes, and surface area in five of the six diameter range classes isometrically scaled to both above- and below-ground biomass (Figure 3A). Both root length and surface area in diameter class 1 appeared to take on an intermediate value between 0 and 1 for both above- and below-ground biomass, which would suggest a faster accumulation of biomass relative to either of these traits. We did not observe any traits to be isometric to variation that was unique to above- or below-ground biomass, but there were relationships that appeared to be distinct from both 0 (no relationship) and 1 (isometry), representing negative allometries. No proportion-based traits were found to scale isometrically with any feature of plant size. Interestingly, none of the above-ground multidimensional predictor traits captured from the CropReporter, LinkSquare spectrometers, or elemental composition analysis were isometric to any feature of plant size (Appendix S9). This suggested that while root traits being modelled by the multidimensional phenotypes may need to be adjusted for plant size, the multidimensional predictors do not.

**Figure 3:**
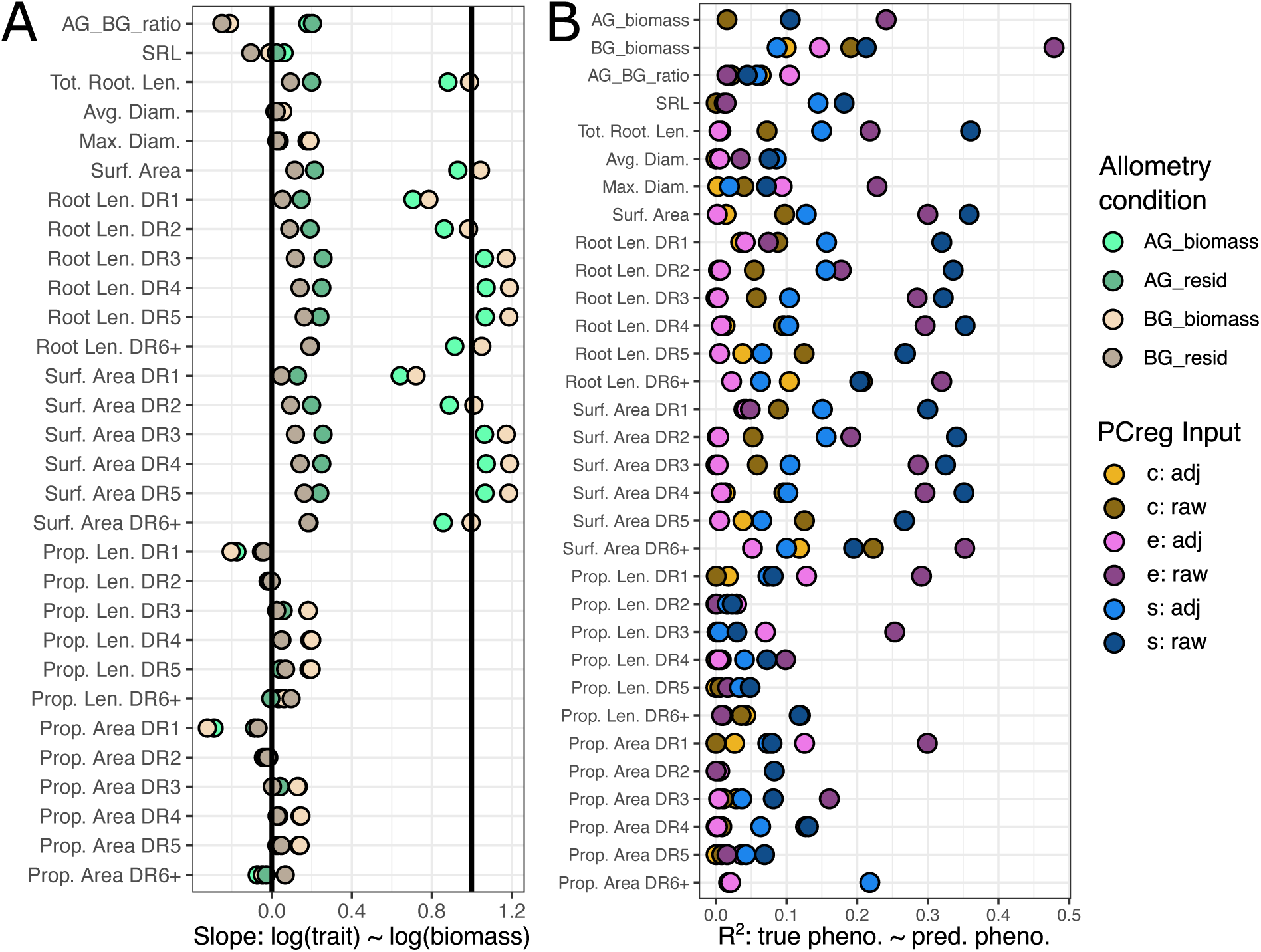
Allometry analysis and multivariate models. (A) The slope of allometric relationships between root traits and four different aspects of plant size: above-ground biomass, below-ground biomass, variation unique to above-ground biomass (AG resid), and variation unique to below-ground biomass (BG resid). Bold vertical lines demonstrate slopes equal to 0 (no relationship) and 1 (isometry or equal scaling). (B) Performance on multivariate predictive models from CropReporter (c), spectral reflectance (s), and elemental composition (e) principal components. Each model was fit with raw values (darker color tones) and after being adjusted (adj) for above-ground biomass (lighter color tones).

### Multivariate predictive models

For each trait, we fit principal component regressions using PCs derived from the CropReporter (3), LinkSquare spectrometers (31) and elemental composition (5; Figure 3B). In general, we found that models parameterized with either spectral or elemental composition tended to perform better (CropReporter vs elemental composition: P < 0.01; CropReporter vs spectra: P < 0.01; elemental composition vs spectra: ns). For example, elemental composition PCs predicted below-ground biomass with an R^2^ = 0.47 (P < 0.01) and spectral PCs predicted total root length with an R^2^ = 0.36 (P < 0.01). Of note, in almost every root trait modelled, adjusting for above-ground biomass worsened the performance of the principal component regressions. Taking the examples above, elemental PCs predicted adjusted below-ground biomass with an R^2^ = 0.15 (P < 0.01) and spectral PCs predicted adjusted total root length with an R^2^ = 0.15 (p < 0.01). Interestingly, in almost every root trait, after correcting for above-ground biomass, spectral PCs were better predictors than elemental composition PCs (CropReporter vs spectra: P < 0.01; elemental composition vs spectra: P = 0.02). Traits derived from proportions were not convincingly predictable in any diameter class by any of the PC modalities, with or without adjustment for above-ground biomass.

While our current capacity for prediction is limited, we note a convincing degree of explanatory power from spectral principal component models. For example, when we use all samples to regress root length and surface area traits across diameter range classes, we see R^2^ values between 0.625 and 0.675 in raw and SNV-transformed spectra (Appendix S3). Moreover, we find that this explanatory power might be tunable using optimization techniques. For example, when we look at total surface area, additional information was gained by incorporating more principal components into the regression (Appendix S10). By estimating the true R^2^ via bootstrapping, we find that a model with 10 PCs has an average R^2^ of 0.41 while a model with 37 PCs has an average R^2^ of 0.90. We caution over-interpretation of these models much past this as the models become over-parameterized, but the information gained by additional PCs suggests there is still room for considerable model improvement.

### Leaf spectral associations with root traits

Because we demonstrated that spectra obtained from leaves hold some predictive capacity for below-ground traits, even after accounting for above-ground biomass, we sought to understand which regions of spectral reflectance may prove useful in predicting below-ground traits. To do this, we fit wavelength-by-wavelength linear models regressing each trait against each SNV-transformed wavelength (Figure 4). Each wavelength was fit independently for each sensor in the spectrometer and models were fit to both the scaled trait values and traits adjusted for above-ground biomass. We found that root traits correlated with reflectance across the range of spectra analyzed. For example, below-ground biomass, total root length, and surface area saw peaks in the F-statistic around 425 nm, 475 nm, 550 nm, and 650 nm as captured by LinkSquare light source 0. In each case, the scaled trait had a stronger relationship with reflectance than the adjusted trait, but the smoothed average for both forms of the model tended to be significant. Each of these traits were also associated with reflectance around 900 nm. This effect was most strongly captured on LinkSquare light source 0, but was weakly detected by LinkSquare lightsource 1. The ratio between above- and below-ground biomass saw a subset of these peaks, namely at 475, 650, and 740 nm. We also noticed a signal at 850 nm, but this signal was most strong in the adjusted trait models. Interestingly, a large peak in average diameter was detected near 750 nm, but only in the adjusted trait model. Finally, average diameter was associated with reflectance at many wavelengths across light sources and trait adjustments, but these tended to be in very narrow bands, not sweeping areas of the spectrum (Figure 4).

**Figure 4:**
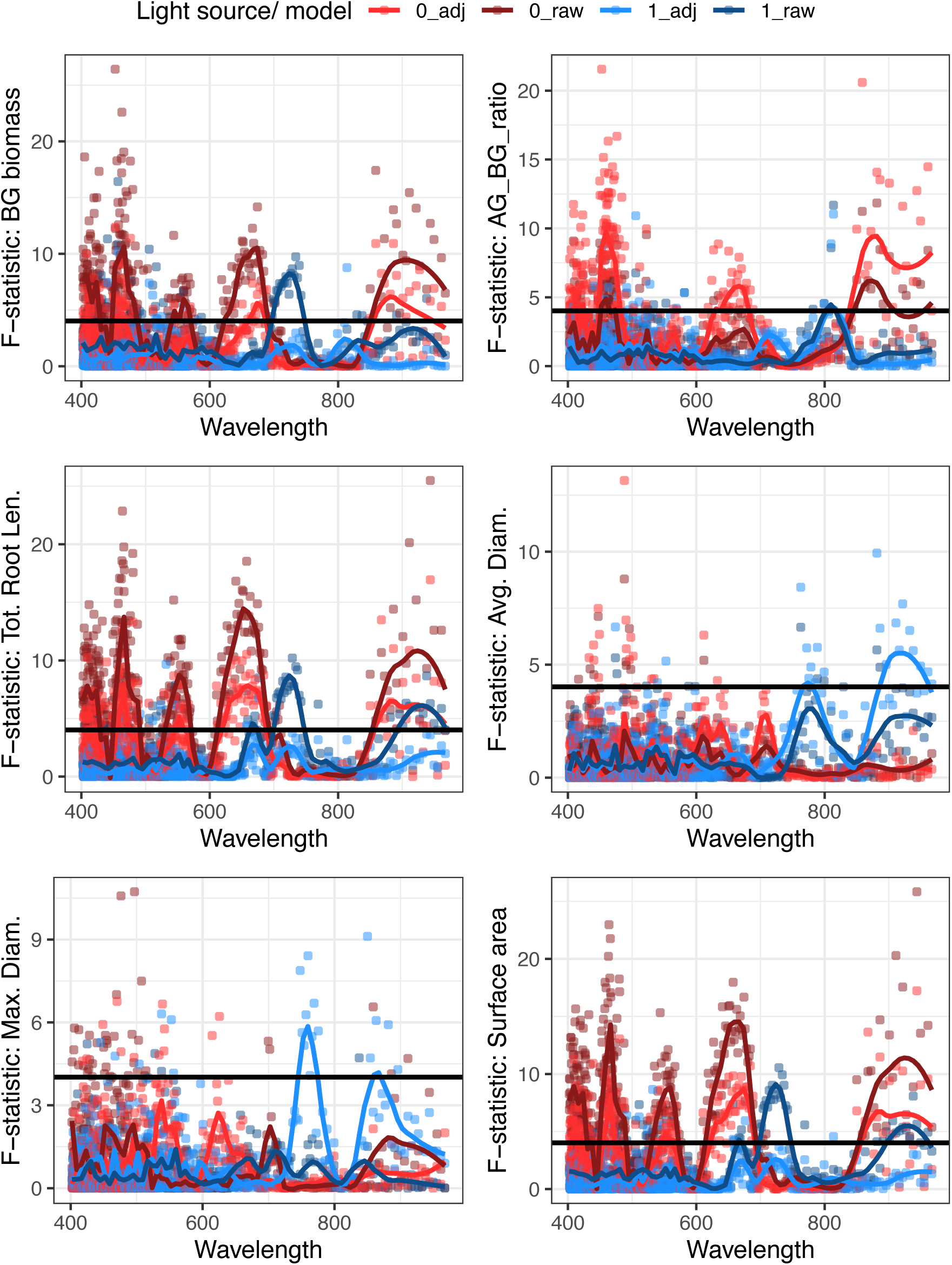
Wavelength-by-wavelength linear models. F-statistics showing the general relationships between reflectance at each wavelength and the trait listed on the y-axis. Traits are z-score transformed. Models were fit independently for each sensor in the LinkSquare. Each model was fit with raw values (darker color tones) and after being adjusted (adj) for above-ground biomass (lighter color tones).

## DISCUSSION

In this study, we explored variation in above- and below-ground phenotypes in American licorice as a function of plant size and across source populations. We demonstrated that both leaf spectral reflectance and leaf elemental composition are reasonable multidimensional phenotypes to survey as proxies for root phenotypes and to use in building robust models for trait estimation. Establishing patterns of correlation between easy-to-measure shoot traits and difficult-to-measure root traits offers one way to assess variation in the hidden half of plants. This approach opens opportunities for evolutionary and ecological analysis of root traits, and holds promise for future breeding pipelines where large populations can be quickly surveyed for root traits.

### Root trait variation across source populations: a foundation for crop improvement and domestication

We characterized the degree to which above- and below-ground phenotypes of American licorice varied across source populations. While most above-ground phenotypes were not significantly variable across source populations, root phenotypes varied across source populations (Figure 1). For example, many aspects of root size including below-ground biomass, SRL, total root length, surface area, length and surface area in the narrowest diameter classes, and the proportions of length and area at the widest diameter class were variable across populations. It is unfortunate that we do not have detailed information on the original location of the source populations; however, we know that seeds were obtained from different climatic regions of the US, including Arizona, Oregon, and across the Midwest (Appendix S1). Roots of these plants would be experiencing vastly different edaphic conditions, therefore, the differences in root phenotypes that we observed are expected. One important limitation of this study is that it was carried out entirely in pots, and previous studies have demonstrated the effects of pot size on root phenotypes (Poorter et al., 2012a). In addition, phenotypic variation described in this manuscript offers only a snapshot of the total intraspecific variation in American licorice. Future work based on field-grown individuals, and involving root chemical analysis, is required for a more comprehensive phenotypic understanding of this taxon.

Variation is the foundation of evolution and is required for breeding (Swarup et al., 2021). Variation in key root traits across populations suggests that breeding programs could select for root system traits that enhance crop performance and resilience, such as increased root mass for nutrient uptake or deeper roots for drought resistance (Lynch, 1995, 2019; Comas et al., 2013). In addition, variation in traits related to root length and architecture may directly affect ecosystem services (Den Herder et al., 2010). The ability to easily assay root trait diversity could accelerate improvement of licorice and other crops selected for their below-ground attributes, leading to cultivars that are targeted for specific agricultural goals or better adapted to specific environmental conditions. A key part of this process will be establishing mechanistic links between variation in root traits and desired crop improvement goals in both the root system (e.g., enhanced chemical composition of roots in plants like licorice, and more broadly, ecosystem services provided by root systems) and shoot system (increased yield). In order to hone selection for below-ground traits, it will be necessary to develop a better understanding of the genetic basis of observed variation in root traits, and to characterize variation under field conditions over the course of multiple growing seasons.

### Correlations between above- and below-ground traits suggest potential for enhancing root systems via shoot system traits

One of the most persistent patterns identified in this study is that licorice root traits are correlated with above-ground biomass, suggesting that licorice root traits can likely be assayed by measuring above-ground traits. This may be a unique feature of some systems as resource allocation may favor either root or shoot growth depending on development, environmental conditions, or phylogeny (Brouwer, 1962; Bloom, 1985; Poorter et al., 2012b; Ma et al., 2021).

Every root trait measured in this study was correlated with one or more multidimensional above-ground predictor variables. Most notably, we observed correlations between root traits and leaf spectral reflectance and leaf elemental composition. The observation that leaf elemental composition may correlate with root phenotypic variation is unsurprising given that most elements are taken in via the roots. Previous work has demonstrated that elemental accumulation in the leaves correlated with root length accumulation across depths, likely owing (in part) to unique elemental signatures at different soil depths (Hanlon et al., 2024). Leaf spectral correlates for below-ground traits are more surprising. While broad band indices, like NDVI (Huang et al., 2021), have been shown to correlate with above-ground plant size, the cPCs (principal components derived from CropReporter data) correlated with root traits but not plant size (Figure 2B). Narrow-band spectral measurements were split between PCs that correlated with above-but not below-ground traits (sPC17), those that correlated with above-ground biomass and below-ground traits (sPC2), and those that only correlated with root traits (sPC1, sCP10, sPC24). While leaf spectra have been used to model leaf traits (Kothari et al., 2023), predicting what is happening below ground has proven more challenging. One example is the prediction of potato yield from broadband indices, but the performance of these models are often exceptionally variable (Mukiibi et al., 2024). Models derived from hyperspectral reflectance can perform better (Li et al., 2020), but (like the models reported in this paper) suffer from a tradeoff of sample size and model complexity.

### Predictive models using leaf spectral and elemental data show promise for root trait estimation but require optimization for greater accuracy

The question of how traits correlate with above-ground biomass (or, more generally, plant size) is important as we could approach this study with either of two potential goals: 1) predicting below-ground traits independent of their known correlations or 2) trying to identify above-ground signatures that are unique to each below-ground trait by accounting for known correlates. To formalize the analysis of correlations between below-ground traits and different traits representing plant size, we tested for trait allometry (Niklas, 2004) against four components of plant size: above-ground biomass, below-ground biomass, size unique to the above-ground, and size unique to below-ground. We found that several, though not all, root traits measured in this study are isometric to (scale with) both above- and below-ground biomass, in particular, traits related to root length and surface area (Figure 3A). These traits were all found to be isometric to total size, and not variation unique to either above- or below-ground biomass. We chose above-ground biomass to account for plant size because the allometry results indicated that while both above- and below-ground biomass were isometric to some root traits, there was no unique variation in either trait that scaled with below-ground traits.

In this study, we tested whether multidimensional above-ground phenotypes from the CropReporter, LinkSquare spectrometers, or leaf elemental composition analysis could be used to model below-ground traits (Figure 3B). To account for the isometric relationships with plant size, all models were fit twice: once modeling raw (scaled) trait values and once modeling trait value after adjusting for above-ground biomass. We found that leaf spectral reflectance and elemental composition are the strongest predictors of below-ground traits, but that the models are considerably poorer after adjustment for plant size. Interestingly, after correction for plant size, every below-ground trait except below-ground biomass, the ratio of above- to below-ground biomass, maximum diameter, and some root length proportion traits are better predicted by leaf spectral reflectance than by leaf elemental composition. This, in combination with the observation that the spectral principal component regressions were saturating before all of the spectral variation could be incorporated (Appendix S10), suggests that future work, preferably with a considerably larger sample size, might be able to appropriately model root traits independent of plant size.

To explore the possibility of this assertion, we regressed each of the root traits against wavelength-by-wavelength spectral reflectance. We find that each of the root traits tested (those that were not broken down by diameter range class) were associated with large windows of reflectance, potentially offering insights into which parts of the spectrum may be most informative for predictive modeling. For example, peaks around 425 nm, 475 nm, 550 nm, and 650 nm correspond to regions typically associated with blue, green, and red light. These wavelengths often relate to chlorophyll absorption and reflectance (Gitelson and Merzlyak, 1996; Lichtenthaler and Buschmann, 2001), suggesting that they might be indirectly capturing signals tied to plant vigor or photosynthetic activity, which likely correlate with root traits. Additionally, the strong associations observed around 900 nm, a near-infrared (NIR) region, are of particular interest. NIR reflectance is generally linked to water content, cell structure, and biomass, potentially explaining why traits like below-ground biomass, total root length, and surface area show a strong response here (Peñuelas and Filella, 1998; Clevers et al., 2010).

### Future work

Data presented here demonstrate that observations in American licorice made above ground, including above-ground biomass, leaf spectral signatures, and leaf elemental concentration correlate with root traits. These results are exciting because they suggest it may be possible to gain information about variation in below-ground structures based on data collected above-ground. However, there are limitations of this study that should be considered, as well as additional future work required, in order to more fully develop this approach.

In the case of American licorice or other plants grown for medicinal compounds in their roots, it will be necessary to conduct chemical assays of roots and to link those results with above-ground observations. The primary compound of interest in the cultivation of licorice is the triterpenoid saponin, glycyrrhizin (or glycyrrhizic acid) (PubChem, n.d.; Hayashi et al., 2005), found primarily in its roots. *Glycrrhiza glabra* and glycyrrhizin have rich ethnobotanical histories and are often cited as important components of ‘Traditional Chinese Medicine’ (Ding et al., 2022). In addition to *G. glabra,* other *Glycyrrhiza* species exist on every continent except Antarctica (Duan et al., 2020), with variable chemical compositions and ethnobotanical histories. American licorice has attracted attention for centuries due in part to medicinal properties found in roots, the main storage site for glycyrrhizin and other potentially useful saponin compounds (Hayashi et al., 2005). Although characterization of chemical composition of licorice roots was beyond the scope of this study, based on the variation in root phenotypes observed, we expect variation in chemical composition and/or concentrations (Le Li, Jianping Yong, Canzhong Lu, Danian Tian, 2024).

There is still much to explore in root trait variation, both in species like American licorice where root chemical attributes are the primary targets of selection (Jiang et al., 2020; Wang et al., 2020; Le Li, Jianping Yong, Canzhong Lu, Danian Tian, 2024), as well as in other perennial herbaceous species where roots are valued for their contributions to seed yield and ecosystem services. Additional work is required to understand which root traits are potential targets of selection, and how variation in those specific root traits relate to assayable above-ground features of the plant. It will be important to continue to advance understanding of how aspects of the root system correlate with crop yields, as well as how they contribute important ecosystem services (e.g., carbon sequestration) below ground. With this information in hand, it will be possible to hone predictive approaches based on above-ground observations to enhance and optimize specific aspects of the root system.

## CONCLUSIONS

This study highlights the potential for leveraging spectral reflectance and elemental composition to predict difficult-to-measure below-ground root traits in American licorice. By correlating above-ground phenotypes with root system traits, we demonstrate non-invasive, non-destructive scalable methods to estimate root characteristics. The strong correlations observed between leaf spectral data and root traits suggest that these methods can serve as valuable tools to assay root systems. However, the complexity of plant size isometry and its effect on predictive modeling highlights the need for careful adjustment of plant size in these analyses. Larger, more comprehensive studies are required to refine these models, as work presented here indicates limitations in predictive accuracy, particularly after adjusting for above-ground biomass. Increasing sample size and incorporating more leaf spectral variation could help develop more robust models that better capture the unique variation in root traits. This line of research holds promise as a novel means to capture information about variation in root traits, potentially facilitating evolutionary and ecological analyses of below-ground structures, as well as breeding efforts to select for traits that improve crop performance, resilience, and adaptability.

## Acknowledgements

We thank the Donald Danforth Plant Science Center, Foundation for Food and Agriculture (CA20-SS-0000000123), Herb Society of America, National Science Foundation (BII 2120153), The Perennial Agriculture Project in conjunction with the Malone Family Land Preservation Foundation and The Land Institute, Saint Louis University, and Taylor Geospatial Institute Postdoctoral Fellowship to ZNH for funding this work. We also thank Dr. Arvid Boe for access to seeds, members of the Miller Lab, specifically Katherine Korein, Puja Patel, Jackson Braley, William Keaggy, Zoraya Piedra, Samantha Selby and Felix Vatman, for assistance with data collection, and Danforth Center Plant Growth Facilities (RRID:SCR_024902) and Danforth Center Phenotyping Facility (RRID:SCR_019049).

## Author Contributions

MJR conceived of the experiment and together with AJM provided supervision of project participants. Methodology and investigation were developed and carried out by MJR, VT, MTH, and EP. MJR, ZNH, MTH, and VT contributed to data curation, formal analysis, and visualization of the data. MJR, VT, and ZNH drafted the original manuscript. Grants to MJR and AJM funded the work. AJM administered funding and provided resources to support the work. All authors contributed to reviewing, editing, and revising the manuscript, and all take responsibility for the work described here.

## Data Availability

RhizoVision Explorer output features, CropReporter and associated metadata, spectral reflectance data, elemental composition data, and all R code needed to reproduce the analyses presented in this manuscript can be found on Figshare: 10.6084/m9.figshare.28742870 (Harris, 2025). Raw and segmented root scans are available upon request. Additional supporting information may be found online in the Supporting Information section at the end of the article.

## Online Supporting Information

## Appendix S1

American licorice (*Glycyrrhiza lepidota*) accessions used in this study with accompanying source, domestication status, and location from which it was collected.

| Accession | Source | Status | Location |
| --- | --- | --- | --- |
| 1805 | Chicago Botanic Garden | wild | Kansas, USA |
| 8956 | Chicago Botanic Garden | wild | Nebraska, USA |
| 3232 | Chicago Botanic Garden | wild | Iowa, USA |
| 11_011 | USDA NPGS | wild | Arizona, USA |
| Everwilde Farms | Commercial | improved | - |
| PI215210 | USDA NPGS | wild | Nebraska, USA |
| PI215211 | USDA NPGS | wild | Nebraska, USA |
| PI215212 | USDA NPGS | wild | Nebraska, USA |
| PI215213 | USDA NPGS | wild | Nebraska, USA |
| PI215215 | USDA NPGS | wild | Nebraska, USA |
| Prarie Moon | Commercial | improved | - |
| Sheffields Seed co | Commercial | improved | - |
| W646335 | USDA NPGS | wild | Oregon, USA |
| W649557 | USDA NPGS | wild | Illinois, USA |
| W653059 | USDA NPGS | wild | Illinois, USA |
| Oak Lake | A. Boe | wild | South Dakota, USA |
| RRx22nd | A. Boe | wild | South Dakota, USA |
| SD HWY 18 | A. Boe | wild | South Dakota, USA |
| Felt | A. Boe | wild | South Dakota, USA |

## Appendix S2

Traits loadings onto CropReporter principal components (cPCs)

Appendix S2: Traits loadings onto CropReporter principal components (cPCs)
| Element | PC1 | PC2 | PC3 |
| --- | --- | --- | --- |
| dFv_Fm | 0.15833672 | 0.22488893 | -0.55967 |
| F0 | 0.06146813 | 0.42612809 | 0.22641923 |
| Fm | -0.0577263 | 0.43058379 | 0.1014729 |
| Fv_Fm | -0.1708681 | -0.0953518 | -0.2033505 |
| Red | 0.34739586 | 0.0530839 | 0.0233756 |
| Green | 0.35753306 | 0.03190374 | -0.0056954 |
| Blue | 0.0849279 | 0.29108188 | -0.132894 |
| Hue | -0.2632008 | 0.16921229 | -0.1700927 |
| Saturation | 0.31414137 | -0.1496399 | 0.04486637 |
| Value | 0.35744195 | 0.02800558 | -0.010604 |
| SpcGrn | 0.35465518 | 0.00308587 | 0.01265086 |
| FarRed | 0.24213984 | 0.29297384 | 0.26401292 |
| Nir | -0.0497858 | 0.45602131 | 0.17919815 |
| ChlIdx | -0.3277842 | 0.09262518 | -0.1669411 |
| AriIdx | -0.2700191 | 0.26604893 | 0.16938101 |
| NDVI | -0.0443919 | 0.08879442 | -0.156103 |
| dChl | 0.13400814 | 0.2278756 | -0.5930066 |

## Appendix S3

Visualization of spectral pretreatments and explanatory power of each treatment to predict root traits. (A) Visualization of each spectral pretreatment explored in this study. Pretreatments were applied on a sample-by-sample basis prior to averaging across replicates. For each treatment (Raw, SG, and SNV), we explored the 1st and 2nd derivatives. (B) R-squared from a model regressing each trait against the top 31 spectral PCs from each pretreatment.

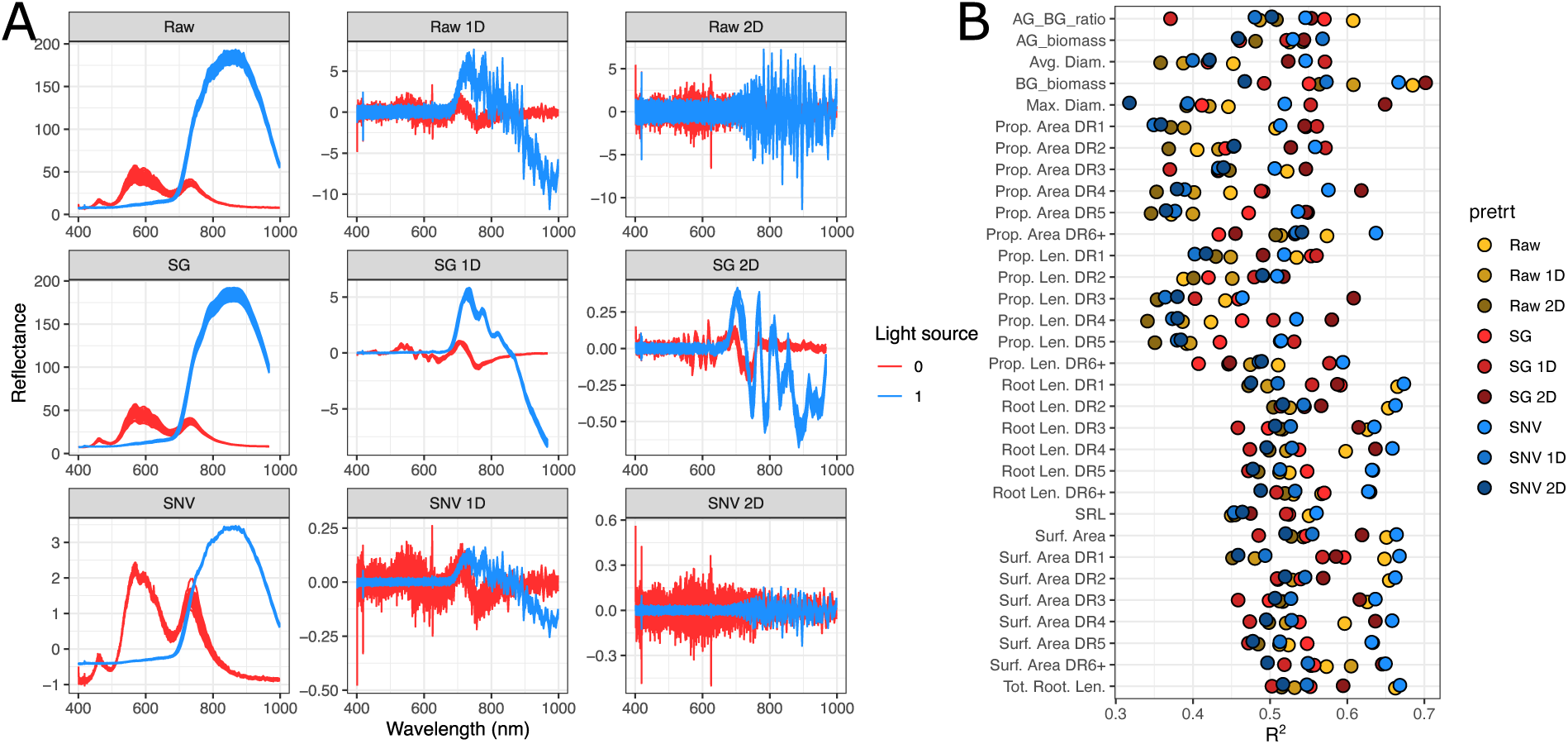

## Appendix S4

Table reporting number of spectral PCs required to explain ∼75% of the total variation in the spectral reflectance across spectral pretreatments and how much variation was explained in each pretreatment after 31 PCs.

| Pretreatment | N. sPCs to Explain 75% | % Variance Explained at 31 PCs |
| --- | --- | --- |
| Raw | 19 | 0.83 |
| Raw1D | 42 | 0.63 |
| Raw2D | 41 | 0.64 |
| SNV | 21 | 0.82 |
| SNV1D | 42 | 0.63 |
| SNV2D | 41 | 0.64 |
| SG | 6 | 0.93 |
| SG1D | 33 | 0.72 |
| SG2D | 40 | 0.64 |
| Mean | 31.7 | 0.72 |

## Appendix S5

Traits loadings onto elemental composition principal components (ePCs)

Appendix S5 Traits loadings onto elemental composition principal components (ePCs)
| Element | ePC1 | ePC2 | ePC3 | ePC4 | ePC5 |
| --- | --- | --- | --- | --- | --- |
| Al27 | -0.1423416 | 0.06735989 | -0.188719 | 0.04837338 | -0.033304 |
| As75 | -0.1572846 | 0.10029285 | -0.3946096 | -0.1012598 | 0.01952123 |
| B11 | -0.3102796 | 0.08301961 | -0.0520619 | 0.02143295 | -0.0891423 |
| Ca44 | -0.3006167 | 0.15744834 | -0.1664113 | 0.04509626 | -0.0098775 |
| Cd111 | -0.2448226 | -0.1138518 | 0.04863003 | -0.2514035 | -0.2759201 |
| Co59 | -0.132557 | -0.0420767 | 0.047134 | 0.70140267 | 0.11340297 |
| Cu63 | -0.2160075 | 0.02806352 | 0.27972281 | -0.0850623 | 0.21159423 |
| Fe54 | -0.1914556 | 0.01265993 | -0.0630791 | 0.45822957 | -0.1151088 |
| K39 | -0.1406829 | -0.5865696 | -0.0396258 | 0.06852349 | 0.13042069 |
| Mg26 | -0.2999292 | 0.20188416 | -0.0676863 | -0.0742061 | -0.0772126 |
| Mn55 | -0.2898937 | 0.00838146 | 0.02268643 | 0.12922308 | -0.0869132 |
| Mo98 | -0.1693064 | -0.0141124 | 0.5140476 | -0.048276 | 0.06419033 |
| Na23 | -0.1950068 | -0.0677894 | -0.2995094 | -0.3819391 | 0.17175718 |
| Ni60 | -0.1017266 | 0.23316094 | 0.4930896 | -0.0849331 | 0.13130048 |
| P31 | -0.1540801 | -0.4730226 | 0.0496467 | -0.0459041 | 0.01274823 |
| Rb85 | -0.2091978 | -0.4402152 | -0.0039999 | -0.0483596 | 0.17327516 |
| S34 | -0.3156485 | 0.06980813 | 0.03222756 | -0.0007652 | 0.03507413 |
| Se78 | -0.2863536 | 0.10939897 | 0.18144751 | -0.146656 | 0.11371809 |
| Sr88 | -0.2885586 | 0.20267609 | -0.1715735 | 0.05591036 | 0.02470991 |
| Zn66 | -0.065587 | -0.1361347 | 0.14771731 | -0.0336361 | -0.8468212 |

## Appendix S6

(trait ranges):

Appendix S6 (trait ranges):
| Trait | Min. | 1st Qu. | Median | Mean | 3rd Qu. | Max. | N |
| --- | --- | --- | --- | --- | --- | --- | --- |
| AG_biomass | 0.27 | 6.1975 | 7.825 | 7.50638889 | 9.21 | 13.33 | 72 |
| BG_biomass | 1.38 | 11.0325 | 16.37 | 15.4732917 | 19.5325 | 30.41 | 72 |
| AG_BG_ratio | 0.19565217 | 0.39655135 | 0.48321716 | 0.53280451 | 0.56234238 | 1.50955414 | 72 |
| SRL | 4.15708237 | 10.1518365 | 14.8295002 | 15.4066625 | 18.9867642 | 30.8898305 | 72 |
| Tot. Root. Len. | 27636.8357 | 135134.443 | 222626.92 | 239425.249 | 326943.32 | 562866.375 | 72 |
| Avg. Diam. | 0.404856 | 0.44694007 | 0.4730353 | 0.48105103 | 0.50460412 | 0.66729648 | 72 |
| Max. Diam. | 10.147645 | 16.0775113 | 17.660024 | 17.4501955 | 19.2450223 | 23.875736 | 72 |
| Surf. Area | 35252.0581 | 188699.149 | 333096.433 | 357977.443 | 488045.622 | 822291.568 | 72 |
| Root Len. DR1 | 7925.71538 | 22689.3384 | 39168.2367 | 41946.0109 | 57596.9107 | 125907.349 | 72 |
| Root Len. DR2 | 12315.794 | 74260.206 | 116993.79 | 128214.174 | 169636.579 | 338596.927 | 72 |
| Root Len. DR3 | 2963.55198 | 20045.4295 | 36230.3665 | 44830.9111 | 67591.9994 | 113010.902 | 72 |
| Root Len. DR4 | 937.046825 | 5982.57855 | 10958.325 | 13371.8706 | 19906.3048 | 38168.3816 | 72 |
| Root Len. DR5 | 329.303782 | 2019.35667 | 3833.86693 | 4684.48397 | 6797.26808 | 13859.5121 | 72 |
| Root Len. DR6+ | 561.951513 | 3468.99862 | 5856.37776 | 6377.79822 | 8641.99615 | 16646.8741 | 72 |
| Surf. Area DR1 | 4829.66263 | 12734.0645 | 20364.6962 | 22149.424 | 30171.79 | 68999.7907 | 72 |
| Surf. Area DR2 | 13309.4685 | 85582.9607 | 132957.644 | 147387.247 | 196377.823 | 387496.938 | 72 |
| Surf. Area DR3 | 5542.37516 | 37398.9766 | 67773.8332 | 83970.9181 | 126763.474 | 212977.226 | 72 |
| Surf. Area DR4 | 2504.97239 | 16015.7456 | 29343.1295 | 35754.8048 | 53161.6791 | 102204.312 | 72 |
| Surf. Area DR5 | 1148.5375 | 7010.86966 | 13302.0415 | 16274.9046 | 23535.4273 | 48128.5555 | 72 |
| Surf. Area DR6+ | 5146.25927 | 33662.8163 | 50337.4316 | 52440.1449 | 69504.9604 | 120371.427 | 72 |
| Prop. Len. DR1 | 0.10947198 | 0.15360276 | 0.17725022 | 0.18632119 | 0.21598706 | 0.32369922 | 72 |
| Prop. Len. DR2 | 0.40120945 | 0.50796883 | 0.53724479 | 0.53252295 | 0.55935851 | 0.62595301 | 72 |
| Prop. Len. DR3 | 0.10723196 | 0.14930988 | 0.17474333 | 0.17806512 | 0.2036067 | 0.25810616 | 72 |
| Prop. Len. DR4 | 0.0280253 | 0.04205025 | 0.04959339 | 0.05396766 | 0.06230261 | 0.11507048 | 72 |
| Prop. Len. DR5 | 0.00877432 | 0.01325028 | 0.01846313 | 0.0194363 | 0.02263773 | 0.05419395 | 72 |
| Prop. Len. DR6+ | 0.01112678 | 0.01968163 | 0.02657747 | 0.02968679 | 0.03592948 | 0.09012357 | 72 |
| Prop. Area DR1 | 0.02641656 | 0.05234288 | 0.06192092 | 0.0682823 | 0.08370033 | 0.13700371 | 72 |
| Prop. Area DR2 | 0.24416432 | 0.36669406 | 0.41204911 | 0.40818966 | 0.4465088 | 0.50827192 | 72 |
| Prop. Area DR3 | 0.13603794 | 0.18949714 | 0.22249922 | 0.22068196 | 0.2485596 | 0.31723279 | 72 |
| Prop. Area DR4 | 0.05705982 | 0.07993172 | 0.09054318 | 0.09415234 | 0.10534499 | 0.16578948 | 72 |
| Prop. Area DR5 | 0.0229391 | 0.03258823 | 0.04122165 | 0.04353582 | 0.04987658 | 0.10095222 | 72 |
| Prop. Area DR6+ | 0.06361393 | 0.11600463 | 0.15321733 | 0.16515792 | 0.20608578 | 0.39166217 | 72 |
| cPC1 | -4.7163905 | -1.9821236 | -0.4102035 | 1.31E-15 | 1.72464936 | 8.27054655 | 72 |
| cPC2 | -4.2371658 | -1.2550434 | -0.0841612 | -1.19E-15 | 1.36684035 | 4.43072156 | 72 |
| cPC3 | -3.076139 | -0.8025462 | -0.0403861 | 5.81E-16 | 0.90002051 | 3.39634186 | 72 |
| sPC1 | -36.681372 | -17.230452 | -1.6610561 | -6.73E-15 | 15.2304666 | 59.0545282 | 73 |
| sPC2 | -25.315783 | -5.3488096 | 0.04636806 | -3.51E-15 | 5.00384503 | 38.4421855 | 73 |
| sPC3 | -14.965071 | -6.8833049 | 0.24988226 | -7.60E-15 | 4.93050308 | 16.4362506 | 73 |
| sPC4 | -15.726637 | -4.7542506 | -0.1479843 | -7.60E-15 | 4.10140196 | 17.0064071 | 73 |
| sPC5 | -12.420012 | -2.8895316 | -0.1123994 | 7.09E-15 | 3.8513956 | 10.4621185 | 73 |
| sPC6 | -13.321534 | -2.83383 | 0.31092166 | 1.46E-15 | 3.19228777 | 10.9857987 | 73 |
| sPC7 | -12.083254 | -3.0700785 | 0.13982449 | -4.14E-16 | 3.26436856 | 14.0625066 | 73 |
| sPC8 | -11.937122 | -1.9390395 | 0.1622901 | -1.20E-15 | 2.39150804 | 11.7804933 | 73 |
| sPC9 | -9.3170912 | -2.6796545 | -0.2020101 | 1.98E-17 | 2.48891825 | 10.0151554 | 73 |
| sPC10 | -6.6393379 | -3.1571015 | -0.3559684 | 1.72E-15 | 2.98494844 | 8.10462618 | 73 |
| sPC11 | -9.5185283 | -2.2467804 | 0.42997329 | -3.25E-15 | 2.4211322 | 9.56811428 | 73 |
| sPC12 | -8.898727 | -2.0369532 | 0.21367101 | 6.48E-16 | 2.07010266 | 6.42118319 | 73 |
| sPC13 | -8.2838553 | -2.3202759 | 0.19786006 | 1.30E-15 | 2.33998299 | 7.21032151 | 73 |
| sPC14 | -11.466286 | -1.7660856 | -0.1732127 | -3.07E-15 | 1.43926668 | 7.38691447 | 73 |
| sPC15 | -10.300672 | -1.7387066 | 0.43000779 | 1.02E-15 | 1.44160273 | 5.91906008 | 73 |
| sPC16 | -7.1867307 | -1.6811816 | 0.02959869 | -3.31E-15 | 1.88187294 | 9.3590027 | 73 |
| sPC17 | -6.541455 | -1.4777455 | -0.4273456 | -5.27E-15 | 1.76261313 | 7.20277032 | 73 |
| sPC18 | -6.467581 | -1.6764712 | -0.2862106 | 4.58E-17 | 1.31962409 | 10.0025021 | 73 |
| sPC19 | -8.7462668 | -1.6115433 | 0.30044139 | -2.29E-15 | 1.44056953 | 8.06453402 | 73 |
| sPC20 | -6.0941296 | -1.9574798 | 0.33170957 | -2.74E-15 | 1.46212302 | 5.73602548 | 73 |
| sPC21 | -6.8983145 | -1.9064213 | -0.3097841 | -3.13E-15 | 1.52484693 | 6.55461174 | 73 |
| sPC22 | -7.3340324 | -1.3720708 | -0.0289049 | -4.84E-16 | 1.567399 | 6.78132556 | 73 |
| sPC23 | -7.4099386 | -1.9537178 | -0.5692688 | 2.18E-15 | 1.95682165 | 5.73346055 | 73 |
| sPC24 | -8.1900137 | -1.7151461 | 0.03470296 | -4.38E-15 | 1.76300835 | 6.5499669 | 73 |
| sPC25 | -6.4078307 | -1.7588119 | -0.0931573 | -4.84E-15 | 1.78080575 | 6.64816468 | 73 |
| sPC26 | -7.2967512 | -1.6926631 | 0.33372115 | -2.78E-15 | 1.98105318 | 6.14747218 | 73 |
| sPC27 | -6.8072585 | -1.365935 | 0.10178447 | -9.74E-16 | 1.54688345 | 6.27752575 | 73 |
| sPC28 | -6.9252754 | -1.3209568 | -0.4652061 | 3.75E-15 | 1.65625518 | 6.14737401 | 73 |
| sPC29 | -5.8914104 | -1.9026697 | 0.00427459 | -1.21E-15 | 1.73551788 | 6.76220765 | 73 |
| sPC30 | -5.3541024 | -1.4215151 | 0.12857375 | 6.46E-16 | 1.63243737 | 6.56161063 | 73 |
| sPC31 | -5.0105831 | -2.0211863 | -0.2493401 | -1.80E-15 | 1.57097196 | 7.99541755 | 73 |
| ePC1 | -9.5261679 | -1.5682702 | 0.63029672 | -4.12E-16 | 1.93718841 | 6.2331312 | 68 |
| ePC2 | -4.351008 | -0.6572558 | 0.17123849 | -4.58E-16 | 0.91807396 | 3.03563326 | 68 |
| ePC3 | -2.2315629 | -0.6220446 | -0.2518379 | 4.92E-17 | 0.41340019 | 3.29337744 | 68 |
| ePC4 | -1.9513925 | -0.539185 | -0.1334666 | -4.15E-17 | 0.20914665 | 4.93730126 | 68 |
| ePC5 | -6.719593 | -0.220218 | 0.08418844 | 2.07E-16 | 0.47625093 | 2.13651455 | 68 |

## Appendix S7

Spectral loadings on sPCs. Following selection of the best pretreament, we plot the loading of each wavelength, by sensor, onto each of the top 31 sPCs.

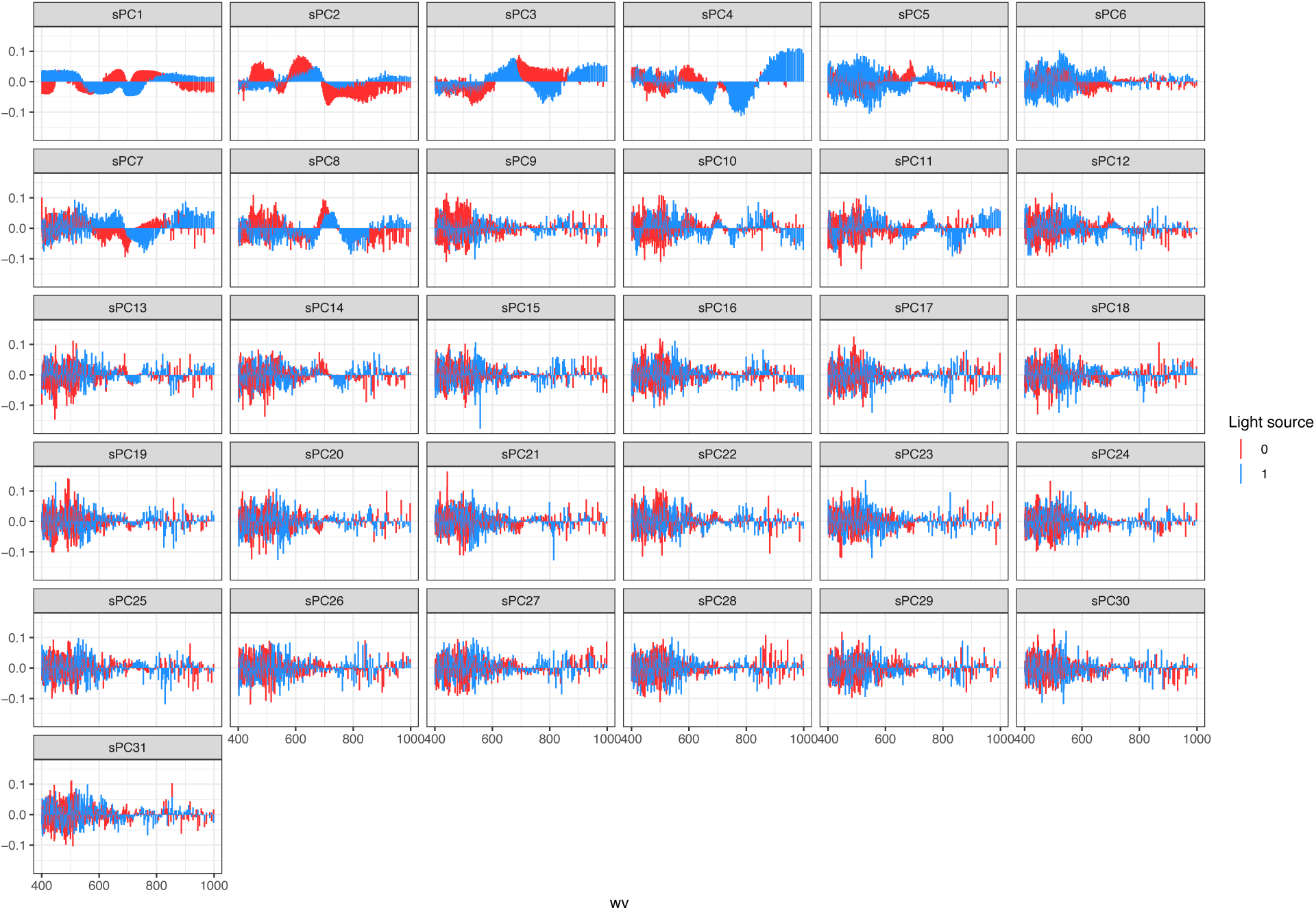

## Appendix S8

Pairwise trait correlations with only the above-ground traits (See Figure 2B).

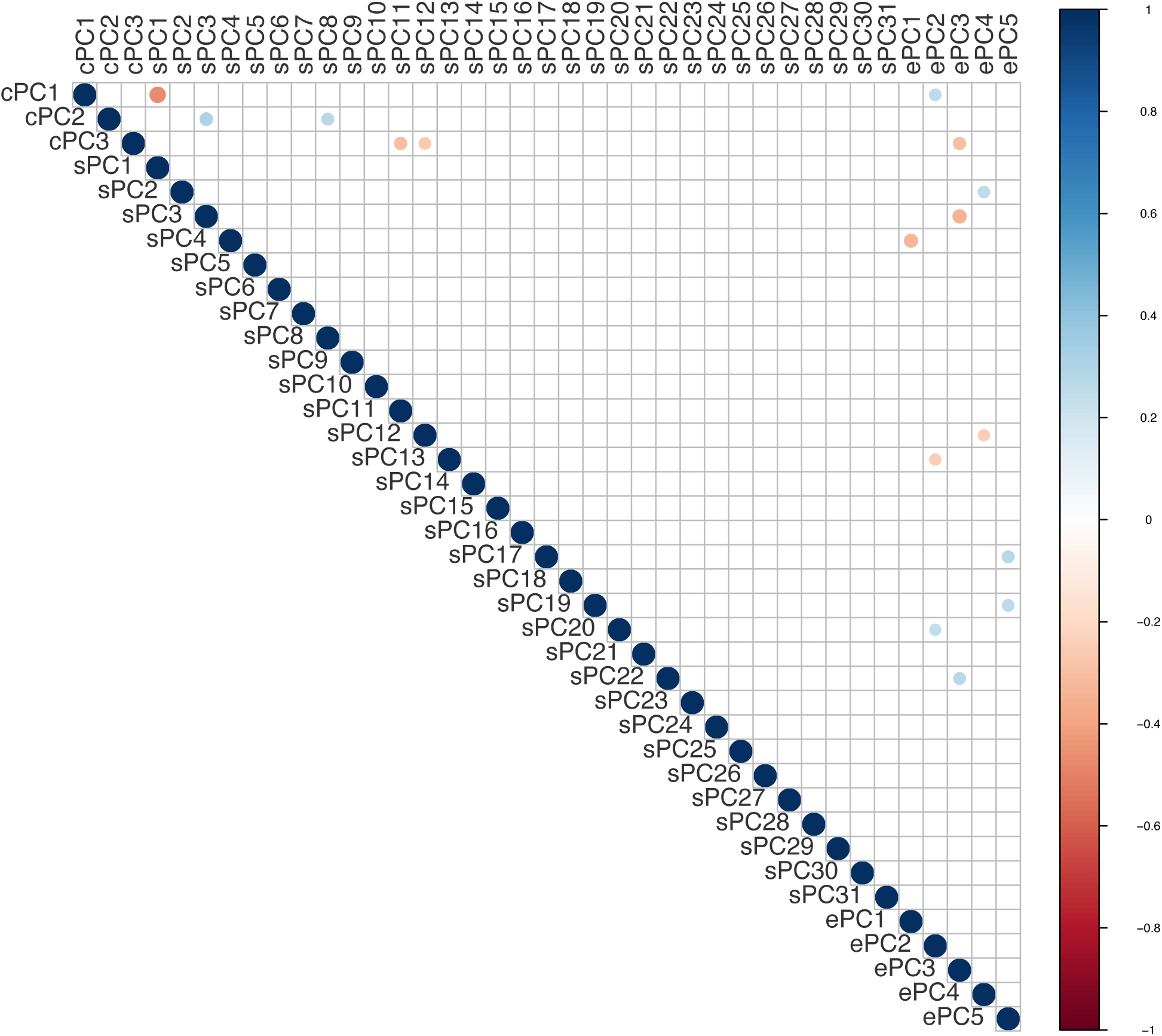

## Appendix S9

Allometric analyses for each of the above-ground traits. See Figure 3A.

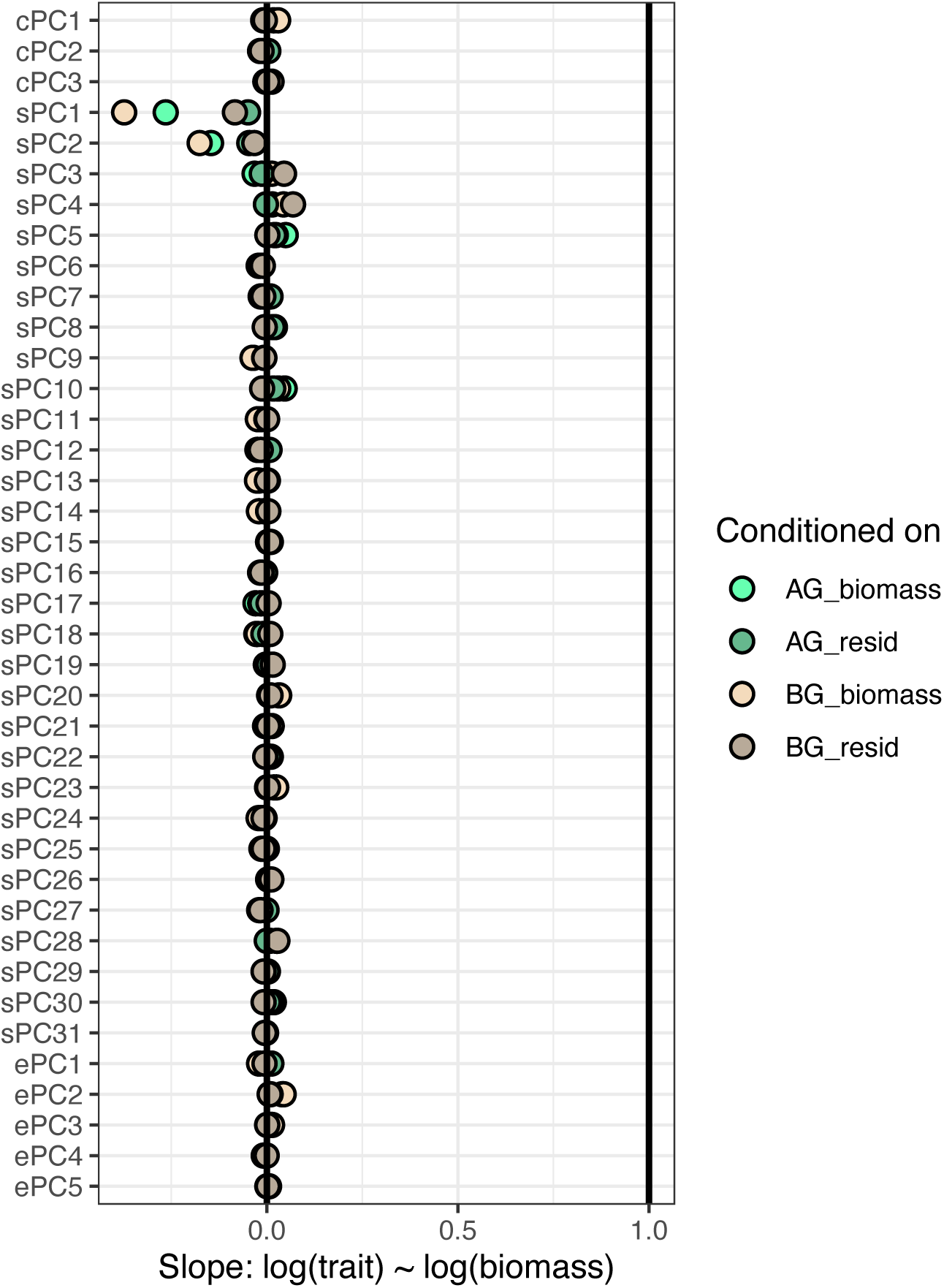

## Appendix S10

Bootstrap analysis for the number of PCs to consider in the best spectral pretreatment. The red vertical line shows the number selected for this study (31).

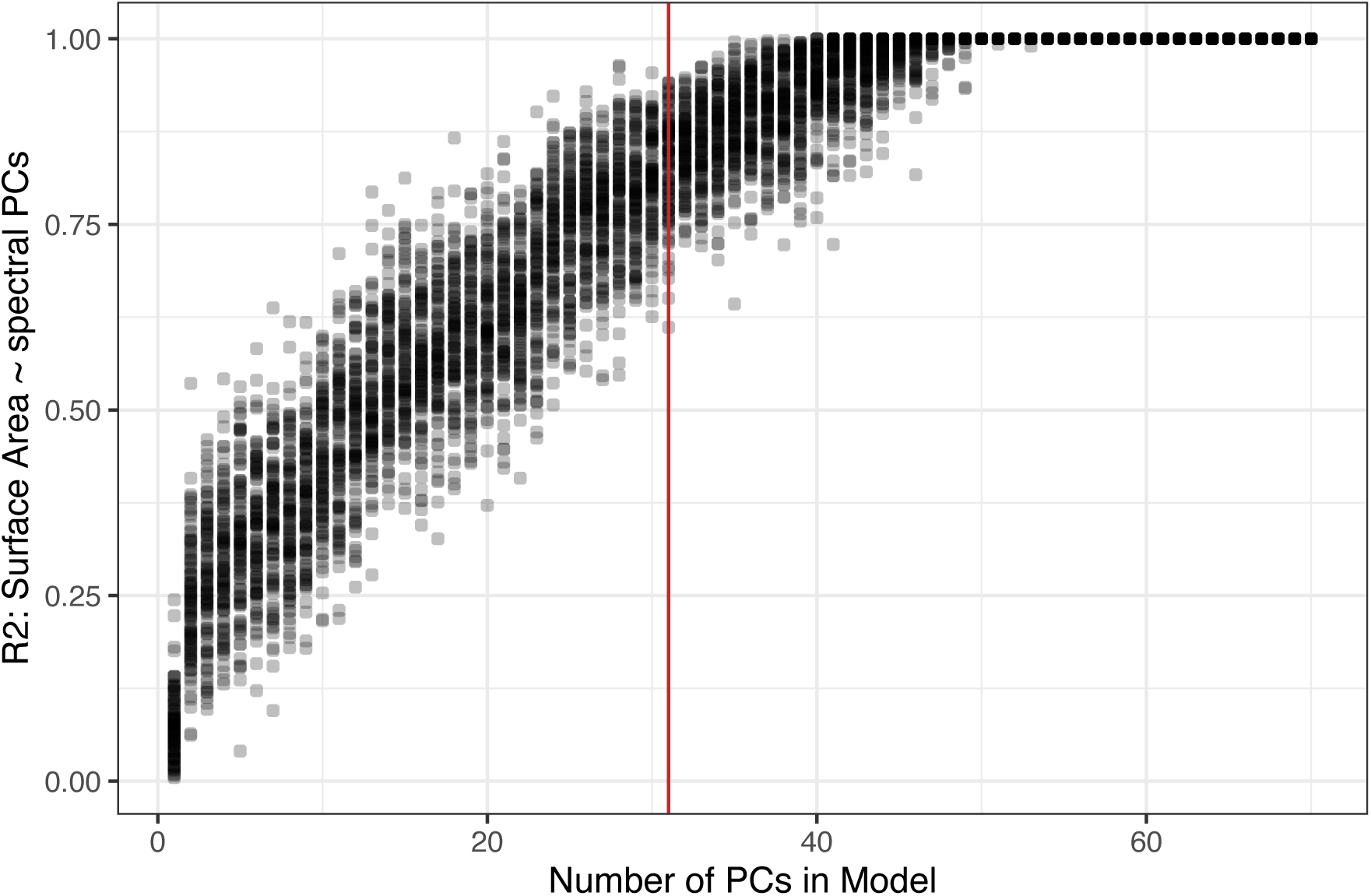

